# Combining LSD1 and JAK-STAT inhibition targets Down syndrome-associated myeloid leukemia at its core

**DOI:** 10.1101/2022.01.14.476342

**Authors:** Juliane Grimm, Raj Bahyadia, Lucie Gack, Dirk Heckl, Jan-Henning Klusmann

## Abstract

Children with Down syndrome (DS) are predisposed to developing megakaryoblastic leukemia (ML-DS) and often experience severe toxicities from chemotherapy, highlighting the need for targeted therapies with beneficial risk profiles. The genomic landscape of ML-DS is characterized by a combination of mutations in signaling pathway genes and epigenetic modifiers, while aberrant lysine specific demethylase 1 (LSD1) and JAK-STAT activation have both been implicated in leukemogenesis. Here, we demonstrate that combined LSD1 and JAK1/2 inhibition exerts synergistic anti-leukemic effects specifically in ML-DS, both *in vitro* and in patient derived xenografts *in vivo*. The JAK1/2 inhibitor ruxolitinib enhanced the LSD1 inhibitor-induced differentiation, proliferation arrest and apoptosis in patient-derived leukemic blasts. At the transcriptional level, the combination synergistically repressed gene expression signatures essential for cell division. We further observed an immunogenic gene expression pattern in the form of increased cytokine signaling, which – by sensitizing ML-DS blasts to the JAK-STAT signaling blockade induced by ruxolitinib – could explain the increased susceptibility of ML-DS blasts to combination therapy. Taken together, we establish combined LSD1 and JAK-STAT inhibition as an efficacious therapeutic regimen specifically designed to target important steps in ML-DS leukemogenesis, paving the way for targeted therapies in this entity.

## Introduction

Individuals with Down syndrome (DS) are predisposed to developing acute megakaryoblastic leukemia within their first years of life.^1, 2^ Evolution of this myeloid leukemia associated with Down syndrome (ML-DS) occurs in a step-wise process originating from pre-malignant transient abnormal myelopoiesis (TAM) *in utero*.^3–5^ The molecular mechanisms underlying the progression from TAM to overt ML-DS are not fully understood. However, it was previously shown that epigenetic changes play a pivotal role in ML-DS leukemogenesis.^6–8^ The lysine demethylase LSD1 was identified as crucial player in this process, as LSD1-driven gene signatures are highly activated in ML-DS.^8^ LSD1 is essential for normal hematopoiesis, particularly during terminal granulocytic and erythroid differentiation.^9^ Yet, it was also shown to contribute to differentiation blockade in different acute myeloid leukemia (AML) subtypes, *e*.*g. KMT2A*-rearranged leukemias.^10–16^ Upon LSD1 inhibition, increased chromatin accessibility and activation of enhancers and important myeloid transcription factors (*e*.*g*. PU.1 and CEBPα) ultimately induced the differentiation of AML blasts.^13, 17–23^

Consequently, various irreversible LSD1 inhibitors have been developed in recent years, with some currently undergoing clinical trials for AML.^24–27^ To further enhance the anti-leukemic effects of existing treatments, various drug combinations containing LSD1 inhibitors have been studied to date, and shown synergy in the induction of differentiation and cytotoxicity, *e*.*g*. in combination with azacitidine, retinoic acid, mTOR or BET inhibitors.^21, 22, 28–31^

Focusing on ML-DS, the combined inhibition of LSD1 and JAK-STAT signaling seems to be desirable, as acquisition of activating mutations in Janus kinases (JAK) and cytokine receptors are frequently observed in this entity.^6–8^ The combination of LSD1 inhibition and the JAK1 and JAK2 inhibitor ruxolitinib was previously examined in *JAK2*^*V617F*^ mutated myeloproliferative neoplasms, secondary AML and a *CSF3R*^mut^ and *CEBPα*^mut^ AML model, where it showed promising anti-leukemic effects.^18, 20, 32^

Here, we investigated the therapeutic potential of LSD1 inhibition alone or in combination with JAK-STAT signaling blockade in ML-DS – an entity which requires targeted therapies with reduced side effects, as infants with DS often experience severe toxicities when receiving standard chemotherapy.^33, 34^ Thereby, we demonstrate robust myeloid differentiation and proliferation arrest after treatment with the irreversible LSD1 inhibitor T-3775440.^35, 36^ However, combining LSD1 inhibition with ruxolitinib synergistically increased the anti-leukemic effects *in vitro* and *in vivo*, proving the feasibility of a combination therapy specifically adjusted to the mechanisms of ML-DS leukemogenesis.

## Methods

### Cell culture and patient samples

All human cell lines were purchased from the German National Resource Center for Biological Material (DSMZ, Braunschweig, Germany) and cultured as recommended by the supplier. Patient samples (expanded via xenotransplantation) were cultured in StemSpan (StemCell Technologies, Vancouver, Canada) supplemented with streptomycin/penicillin (Thermo Fisher Scientific, Waltham, MA, USA) and a cytokine cocktail (FLT3-L, SCF, IL-6, IL-3, TPO; all purchased from PeproTech, Rocky Hill, NJ, USA). Samples were taken from pediatric AML patients treated as part of the AML Berlin-Frankfurt-Münster study group. Written informed consent was given by all custodians of the participants in consent with the Declaration of Helsinki and the study was approved by the local ethic committees of the participating hospitals.

### Evaluation of drug response and synergy analyses

Cell lines and patient samples were treated for six days. On day 0 and day 3, a serial dilution of T-3775440 (Hycultec, Beutelsbach, Germany) was added to the cells. Medium was supplemented with fresh ruxolitinib (Hycultec) from day 3 to 5. Cell viability was determined on day 6 using the CellTiter-Glo® 2.0 Cell Viability Assay (Promega, Madison, WI, USA) according to the manufacturer’s recommendations. Synergy was analyzed using the SynergyFinder web application (version 2.0).^37^

### Flow cytometry: differentiation, apoptosis and cell cycle assays

All flow cytometry analyses were performed using the CytoFLEX platform (Beckman Coulter, Brea, CA, USA). For assessment of myeloid differentiaton after T-3775440 treatment, anti-CD11b-PC7 (Beckman Coulter) and anti-CD86-Alexa Fluor® 700 (BD Biosciences, Franklin Lakes, NJ, USA) were applied. Evaluation of apoptosis and cell cycle progression was performed using the Annexin V Apoptosis Detection Kit II and the BrdU Flow Kit (BD Biosciences), respectively. Both assays were performed according to manufacturer’s recommendations.

### In vivo experiments

Humanized immunodeficient mice^38^ were bred and maintained at the animal facility of the Martin-Luther-University Halle-Wittenberg. Patient samples (expanded via xenotransplantation) were transplanted via tail vein into sublethally irradiated mice. For monitoring of peripheral engraftment, mice were bled every two weeks starting from six weeks after transplantation. Upon stable engraftment, treatment with T-3775440 (20 mg/kg) and/or ruxolitinib (60 mg/kg) was initiated. Both drugs were administered once daily via oral gavage. After a treatment period of 7 days the experiment was terminated. Leukemic infiltration of the spleen and bone marrow was evaluated by flow cytometry applying a standard antibody panel consisting of anti-mouse CD45-FITC, anti-human CD45-APC, anti-human CD33-PE, anti-human CD3-APC-H7, and anti-human CD34-PerCP-Cy™5.5 (all purchased from BD).

All *in vivo* experiments were approved by local authorities and were performed in accordance to national laws and regulations.

### mRNA sequencing

For total mRNA sequencing analysis, two ML-DS samples (expanded via xenotransplantation) were treated with DMSO, T-3775440, ruxolitinib or the combination of T-3775440 and ruxolitinib for two days. Afterwards, cells were harvested and RNA was isolated using the Quick RNA Miniprep kit (Zymo Research, Irvine, CA, USA) following the supplied protocol. mRNA sequencing, alignment of the RNA reads to GRCh38 genome, and calculation of differentially expressed genes were performed by Novogene (Cambridge, UK). The clustering was performed based on concordant and opposed gene regulation signatures in the monotherapies. Gene set enrichment analysis (GSEA) and gene ontology analysis using the PANTHER classification system were performed as previously described.^39–42^

### Western Blotting

Cells were harvested after treatment with DMSO, T-3775400, ruxolitinib or both drugs in combination and lysed using RIPA buffer (Thermo Fisher Scientific). Protein lysates were separated by SDS-PAGE and blotted onto a PVDF membrane (BioRad, Hercules, CA, USA). Staining was performed using the following antibodies: anti-human STAT3, anti-human phosphor-STAT3 (Y705) (Cell Signaling Technology, Danvers, MA, USA). Imaging was performed using the ChemiDoc Imaging System (BioRad).

### Statistical analyses

For statistical analyses, Graphpad PRISM software (version 9.0.1) and R-Studio (R version 3.6.3)^43^ were used. Two-tailed Student’s t-tests were used for comparisons between two groups. *P*<0.05 was considered statistically significant. All statistical tests and sample numbers are disclosed in the respective figure legends.

## Results

### LSD1 inhibition leads to proliferation arrest and induces differentiation in pediatric acute megakaryoblastic leukemia

RNA-sequencing analysis of different pediatric AML subtypes revealed that LSD1 was highly expressed in pediatric acute megakaryoblastic leukemia (AMKL), and especially in TAM and ML-DS patients (Figure 1A), providing rationale for the use of LSD1 inhibitors in these entities. Consistent with this, the non-DS-AMKL cell line M-07e and the ML-DS cell line CMK were particularly sensitive to irreversible LSD1 inhibition (IC50M-07e = 9.1nM; IC50CMK = 38.8nM; Supplemental Figure 1). Testing serial dilutions of the irreversible LSD1 inhibitor in non-DS-AMKL and ML-DS patient samples expanded via xenotransplantation (see Supplemental Table 1 for patient characteristics), both entities were equally sensitive to LSD1 inhibition in terms of IC50 values (non-DS-AMKL: IC50#1 = 15.0nM, IC50#2 = 2.0nM; ML-DS: IC50#1 = 31.2nM, IC50#2 = 17.1nM, IC50#3 = 3.8nM) compared to normal CD34^+^ hematopoietic stem/progenitor cells. Yet, all dose-response curves plateaued at a certain LSD1 inhibitor concentration (Figure 1B). The non-linear relationship between cytotoxicity and inhibitor dosage points towards proliferation arrest and differentiation in response to LSD1 inhibition rather than cell death. In line with this, we observed myeloid differentiation upon visual inspection (Figure 1C) and upregulation of the myeloid markers CD86 and CD11b after three days of LSD1 inhibitor treatment (Figure 1D).

**Figure 1.**
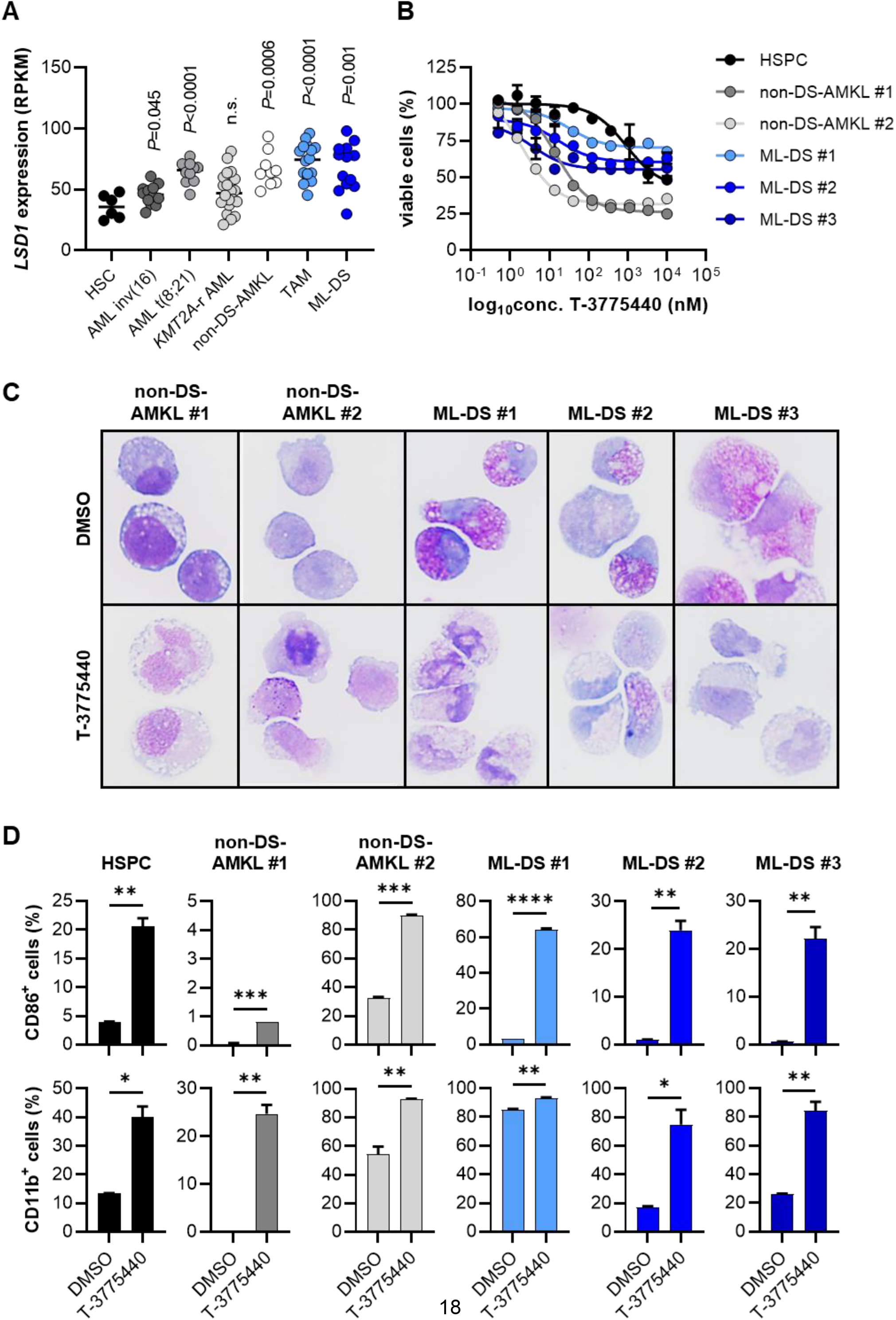
Pharmacological LSD1 inhibition potently induces proliferation arrest and myeloid differentiation in ML-DS and non-DS-AMKL cells. (A) LSD1 expression in different AML subgroups displayed as RPKM (RNA-seq). The *P* values result from pairwise comparisons with normal HSCs. (B) Dose-response curves depicting cell viability of CD34^+^ HSPCs and patient samples (expanded via xenotransplantation) after six days of treatment with serial dilutions of the LSD1 inhibitor T-3775440 for six days. All cell viability values were normalized to the corresponding DMSO control. (C) May–Grünwald–Giemsa stained cytospins of patient blasts treated with DMSO or 350nM T-3775440 for three days. (D) Bar plots depicting cell surface expression of CD86 or CD11b on HSPCs and patient blasts after treatment with DMSO or T-3775440 for three days. Surface marker expression was determined via flow cytometry. \**P*<0.05, \*\**P*<0.01, \*\*\**P*<0.001, \*\*\*\**P*<0.0001; *P* values are derived from two-tailed Student’s t-tests comparing two groups. ML-DS, myeloid leukemia associated with Down syndrome; non-DS-AMKL, acute megakaryoblastic leukemia not associated with Down syndrome; AML, acute myeloid leukemia; RPKM, reads per kilobase of transcript per Million mapped reads; HSC, hematopoietic stem cells; HSPC, hematopoietic stem and progenitor cells; DMSO, dimethyl sulfoxide.

### Inhibition of LSD1 and JAK-STAT signaling acts synergistically to exert anti-leukemic effects in ML-DS

Our initial results revealed a potent block of proliferation and induction of differentiation in non-DS-AMKL and ML-DS samples, however, the therapeutic efficacy of LSD1 inhibition alone may be limited due to its non-linear dose-response relationship. We therefore aimed to design a rational drug combination, such as with the JAK1 and JAK2 inhibitor ruxolitinib, as ML-DS is frequently characterized by autonomous JAK-STAT signaling due to acquired activating mutations.^6–8^ After pre-treatment with 350nM LSD1 inhibitor for three days, cells were treated with serial dilutions of ruxolitinib leading to synergistic growth inhibition in non-DS-AMKL and ML-DS cell lines (Supplemental Figure S2), as well as in all ML-DS patient samples (Figure 2A). The combination of LSD1 inhibition and ruxolitinib also proved to be very effective in non-DS-AMKL blasts, however, with only additive cytotoxic effects in one of the two patient samples (Figure 2A). Drug synergy in the ML-DS samples was also confirmed when calculating the Bliss synergy scores (Figure 2B). Interestingly, samples ML-DS #1 (*JAK1* mutated) and #2 (wild-type for *JAK1, JAK2*, and *JAK3*, Supplemental Figure S3) showed particularly high synergy scores (ML-DS #1 synergy score = 10.4; ML-DS #2 synergy score = 15.6; Figure 2B). On the other hand, the *JAK3* mutated patient sample ML-DS #3 (Supplemental Figure S3) only showed mild drug synergy between LSD1 inhibition and ruxolitinib (synergy score = 2.0; Figure 2B).

**Figure 2.**
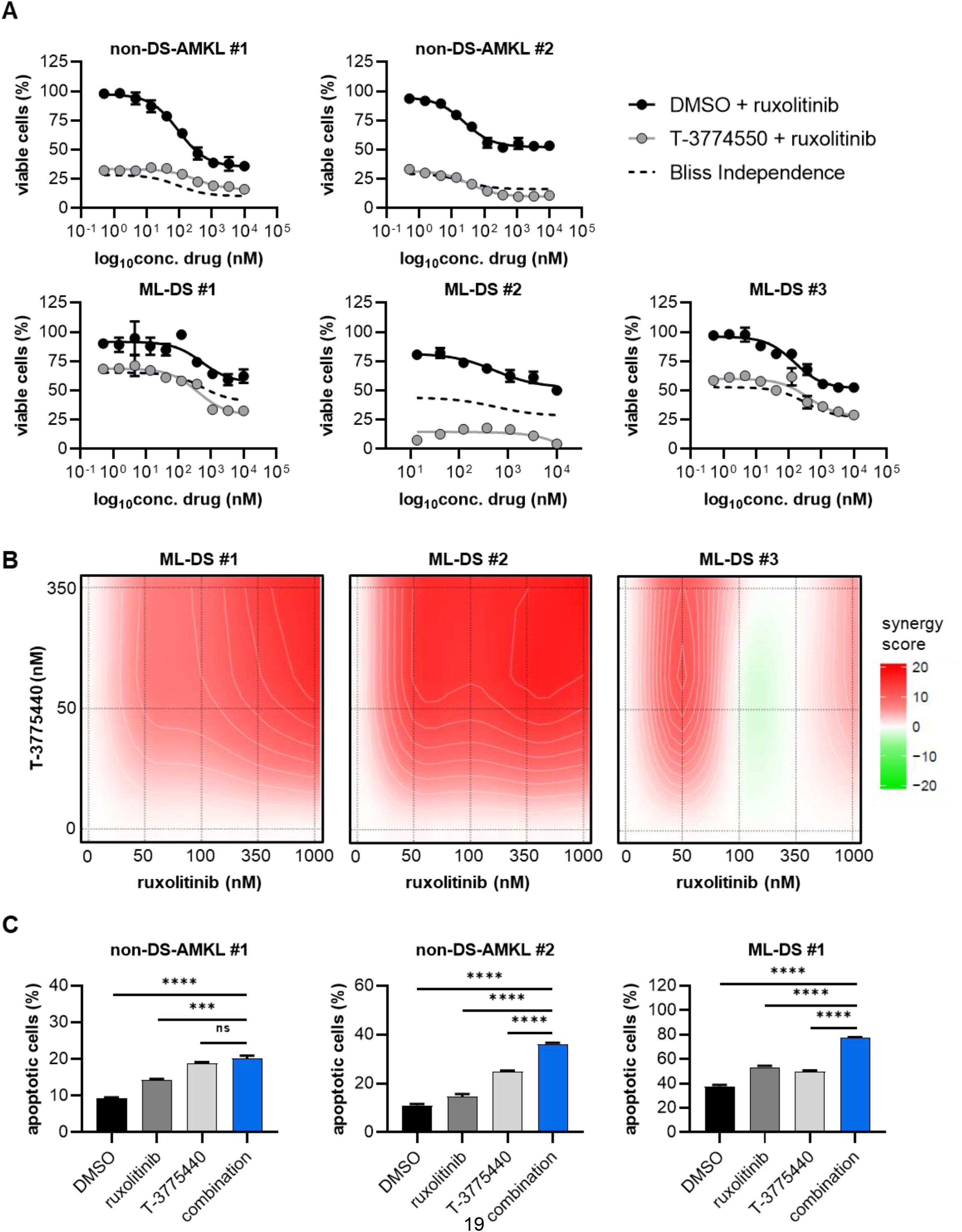
Combined LSD1 inhibition and JAK-STAT signaling blockade synergizes to produce an anti-leukemic effect in ML-DS. (A) Dose-response curves depicting cell viability of different ML-DS and non-DS-AMKL patient samples (expanded via xenotransplantation) after treatment with DMSO or 350nM T-3775440 for six days, with the addition of serial dilutions of ruxolitinib from day 3 to day 6. All cell viability values were normalized to the corresponding all DMSO control. (B) Synergy blots displaying the color-coded Bliss synergy score, calculated after treatment of different ML-DS patient samples (expanded via xenotransplantation) with T-3775440 and ruxolitinib for six days. Interpretation of synergy scores: >10 drug synergy; −10 to 10 additive effects; <-10 drug antagony.^37^ (C) Bar plots depicting percentage of apoptotic cells after treating patient samples (expanded via xenotransplantation) with DMSO, 1µM ruxolitinib, 350nM T-3775440, or the combination of T-3775440 and ruxolitinib for three days. Apoptosis was determined using flow cytometry on DAPI and Annexin V. All Annexin V positive cells were considered apoptotic. ns *P*>0.05, \*\*\**P*<0.001, \*\*\*\**P*<0.0001; *P* values are derived from two-tailed Student’s t-tests comparing two groups. ML-DS, myeloid leukemia associated with Down syndrome; non-DS-AMKL, acute megakaryoblastic leukemia not associated with Down syndrome; DMSO, dimethyl sulfoxide; DAPI, 4′,6-diamidino-2-phenylindole.

Ruxolitinib and the LSD1 inhibitor induced apoptosis to a small extent when used as monotherapies, however, the combination synergistically increased the percentage of apoptotic cells in ML-DS samples and also in one out of two non-DS-AMKL samples (Figure 2C). Moreover, LSD1 inhibition alone or in combination with ruxolitinib effectively blocked G1 to S phase transition (Supplemental Figure S4). This effect was particularly strong in the ML-DS sample.

### Combination of LSD1 inhibition and ruxolitinib synergistically reduce leukemic burden *in vivo*

To further investigate the synergistic anti-leukemic effects of LSD1 inhibition and JAK-STAT blockage *in vivo*, we treated recipient mice with stable engraftment (median peripheral blasts 3.0%, Supplemental Figure S5) of ML-DS blasts (*JAK1* mutated; Figure 3A). At the end of the treatment period, we observed significantly reduced spleen weight in mice treated with ruxolitinib (median 49.2mg) and the combination therapy (median 56.8mg) as opposed to the placebo (median 101.2mg) and LSD1 inhibitor (median 157.0mg) arms (Figure 3B). The spleen infiltration of myeloid blasts was reduced by both monotherapies and the drug combination compared to the placebo group (median percentage of human CD45^+^/CD33^+^ cells: placebo – 28.4%, LSD1 inhibitor – 4.8%, ruxolitinib – 3.9%, combination therapy – 4.2%, Supplemental Figure S6). Of note, the combination of the LSD1 inhibitor and ruxolitinib was the only treatment regimen that achieved a significant reduction of leukemic burden in the bone marrow of the mice (median percentage of human CD45^+^/CD33^+^ cells: placebo – 82.0%, LSD1 inhibitor – 69.5%, ruxolitinib – 85.5%, combination therapy – 55.2%, Figure 3C), underlining the synergistic cytotoxic effect of LSD1 inhibition and JAK-STAT signaling blockade in ML-DS.

**Figure 3.**
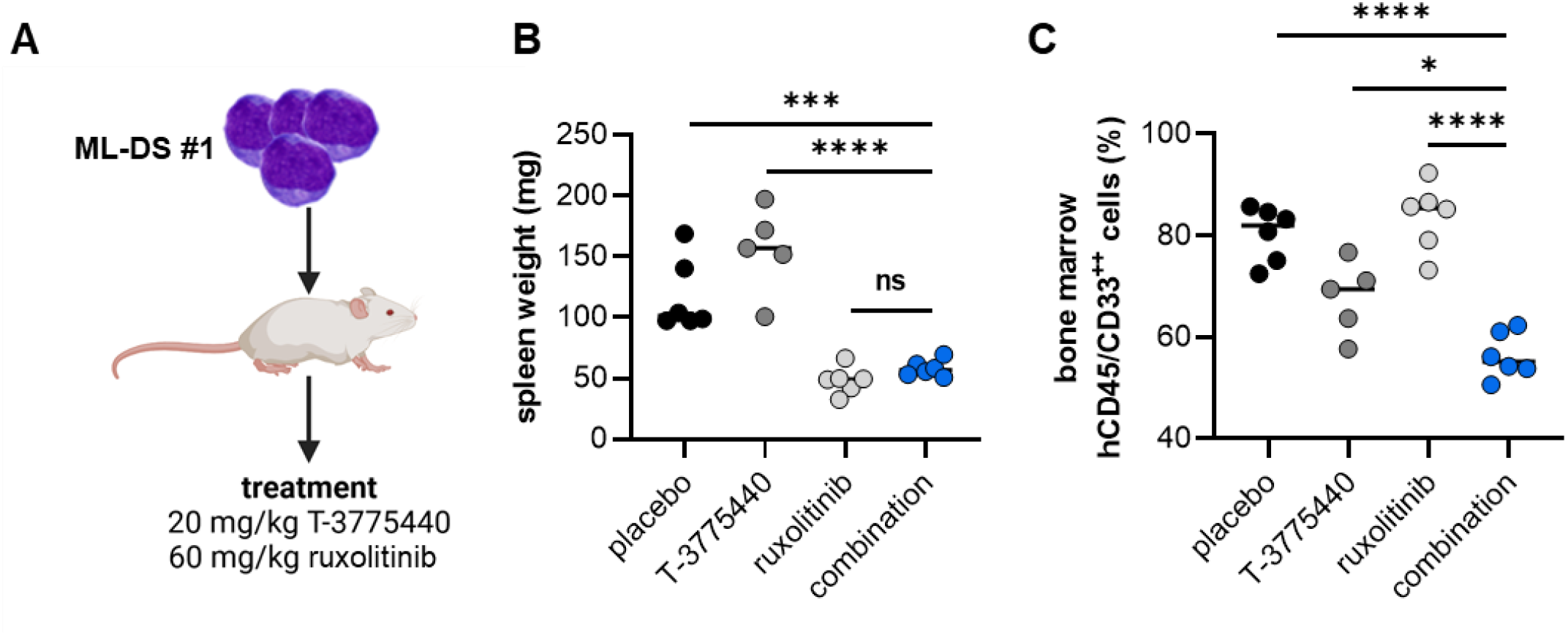
Combined treatment with T-3775440 and ruxolitinib significantly reduces leukemic burden in an *in vivo* model of ML-DS. (A) Treatment schedule of humanized recipient mice^38^ transplanted with the ML-DS #1 patient sample. (B) Spleen weights in milligrams of mice engrafted with the ML-DS #1 patient sample, after treatment with placebo, T-3775440, ruxolitinib, or the combination of both drugs for seven days. (C) Bone marrow infiltration in mice engrafted with the ML-DS #1 patient sample, after treatment with placebo, T 3775440, ruxolitinib, or the combination of both drugs for seven days. Infiltration was defined as percentage of human myeloid blasts and was measured by flow cytometry. Human myeloid blasts were defined as CD45^+^ and CD33^+^. ns *P*>0.05, \**P*<0.05, \*\*\**P*<0.001, \*\*\*\**P*<0.0001; *P* values are derived from two-tailed Student’s t-tests comparing two groups. ML-DS, myeloid leukemia associated with Down syndrome.

### LSD1 inhibition and JAK-STAT signaling blockade synergistically inhibit pro-proliferative gene expression signatures

To unravel the molecular mechanisms behind the synergy between LSD1 inhibition and disruption of JAK-STAT signaling, we performed RNA-sequencing of the patient samples ML-DS #1 and #2 after treatment with DMSO, LSD1 inhibitor, ruxolitinib, or the combination of both drugs. Gene expression was normalized to the DMSO control, identifying 552 differentially expressed genes (*P*<0.05) in both ML-DS samples. In general, the gene expression signature in the combination therapy was mainly driven by LSD1 inhibition, since only one gene was significantly divergently expressed between the LSD1 inhibitor and the combination samples (*SLCO2B1*, Figure 4A). We uncovered four different gene expression clusters, with two clusters displaying cooperative inhibition (cluster 1) and induction (cluster 3) of gene expression by LSD1 inhibition and ruxolitinib (Figure 4A). Gene ontology analyses of the clusters revealed that LSD1 inhibition and ruxolitinib repressed hallmarks of cell division, as genes essential for cell cycle checkpoints, synthesis of DNA, fatty acyl-CoA biosynthesis, and lineloic acid metabolism were downregulated in cluster 1 (Figure 4B). Of note, genes involved in the transition from G1 to S phase were also repressed by the combination therapy, in line with the results from our BrdU assay (Supplemental Figure S4).

**Figure 4.**
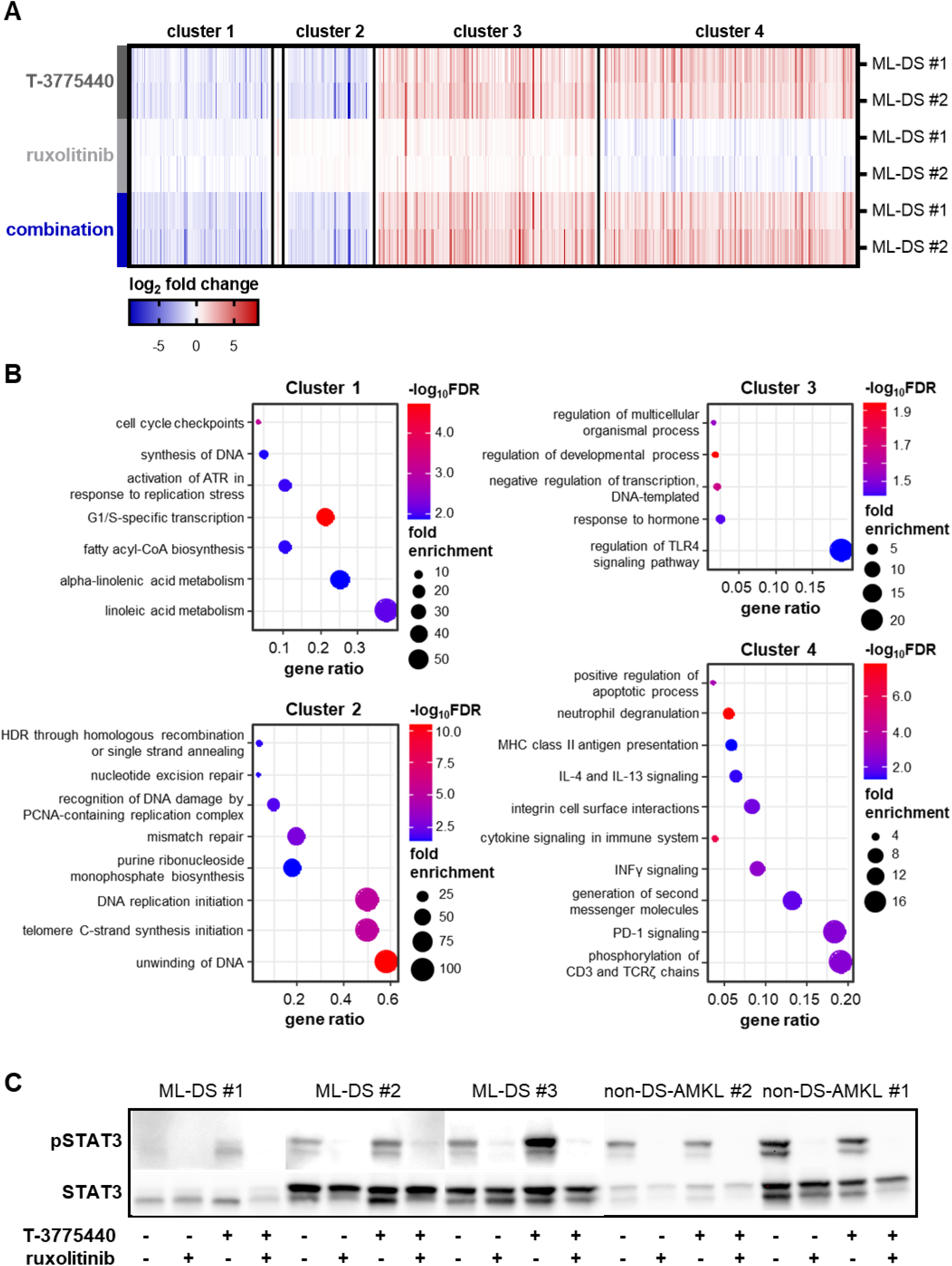
Transcriptomic profiling reveals gene expression signature driven by LSD1 inhibition with induction of cytokine signaling. (A) Heatmap of the 552 differentially expressed genes across all treatment groups (T-3775440, ruxolitinib, combination). Two ML-DS patient samples were used in all groups. Gene expression is normalized to the respective DMSO control and depicted as color-coded log2 fold-change. (B) Gene ontology analysis of the four identified gene clusters from A. The gene ratio was defined as the number of identified genes in a certain biological process, normalized to the total number of genes belonging to this biological process. (C) Western blot of phosphorylated STAT3 and total STAT3 in ML-DS and non-DS-AMKL patient samples (expanded via xenotransplantation) after 3 days of treatment with DMSO, T-3775440, ruxolitinib, or the combination of both drugs. ML-DS, myeloid leukemia associated with Down syndrome; DMSO, dimethyl sulfoxide; non-DS-AMKL, acute megakaryoblastic leukemia not associated with Down syndrome; FDR, false discovery rate.

Independently from ruxolitinib, the LSD1 inhibitor downregulated a plethora of genes involved in DNA replication (cluster 2, Figure 4B). Adding to the anti-proliferative gene signature, ruxolitinib and LSD1 inhibitor cooperatively elevated the expression of negative regulators of DNA transcription (cluster 3, Figure 4B).

### LSD1 inhibition primes cells for ruxolitinib treatment by inducing cytokine signaling in ML-DS

In our *in vitro* studies, we observed that LSD1 inhibition induces myeloid differentiation – a finding that was reflected by the upregulation of genes involved in neutrophil degranulation (cluster 4, Figure 4B). We also demonstrated upregulation of immunological gene expression patterns, e.g. MHC class II antigen presentation, interferon gamma and PD-1 signaling (cluster 4, Figure 4B), upon LSD1 inhibition as monotherapy or in combination with ruxolitinib. Additionally, the expression of gene signatures involved in activation of cytokine signaling was driven by LSD1 inhibition (cluster 4, Figure 4B). Elevated cytokine signaling and the enrichment of pathways supporting differentiation were further validated by gene set enrichment analysis on the entire set of differentially expressed genes (Supplemental Figure S7).

In our study, the induction of cytokine signaling seemed to be limited to the ML-DS context, since we observed increased STAT3 phosphorylation after LSD1 inhibitor treatment in the three ML-DS patient samples but not in the non-DS-AMKL patient samples (Figure 4C). The addition of ruxolitinib after LSD1 pre-treatment completely abrogated STAT3 signaling in all tested ML-DS and non-DS-AMKL samples (Figure 4C). These results suggest that LSD1 inhibition may increase the susceptibility of ML-DS blasts to ruxolitinib by promoting cytokine signaling, which is then abruptly terminated by blockade of the JAK-STAT pathway.

## Discussion

Small molecule inhibitors targeting specific molecular lesions or hallmarks of leukemic transformation have been on the rise for the treatment of AML, especially in subgroups of patients with high treatment-related mortality when receiving intensive chemotherapy.^44–46^ A comparable situation can be found in children with ML-DS, who often experience severe toxicities when treated with chemotherapy. Thus, there is a need for novel therapies designed specifically to target key molecular features of ML-DS, aiming to achieve efficacy at least comparable to chemotherapy while providing a more beneficial risk profile.

Here, we demonstrate a pioneering study combining pharmacological LSD1 inhibition with blockade of JAK-STAT signaling – both of which have previously been shown to be deregulated in ML-DS^8^ – as an efficient treatment strategy in the context of ML-DS. Importantly, we also provide mechanistic insights explaining the particular sensitivity of ML-DS blasts to this drug combination.

In line with investigations in other AML subgroups^11–16^, we observed myeloid differentiation after treatment with the irreversible LSD1 inhibitor T-3775440, accompanied by proliferation arrest but only moderate induction of apoptosis in non-DS-AMKL and ML-DS cells. This was also reflected by a plateauing of the dose-response curves after monotherapy with the LSD1 inhibitor. To maximize anti-leukemic activity, we combined LSD1 inhibition with ruxolitinib, which is a JAK1 and JAK2 inhibitor already approved for the treatment of myeloproliferative neoplasms (MPN). The combination of an irreversible LSD1 inhibitor and ruxolitinib so far has only been tested in the subgroup of post-MPN secondary AML and the rare entity of *CSF3R*/*CEBPα* double mutated AML, where it was shown to be efficient *in vitro* and *in vivo*.^18, 20^ In our study, this drug combination proved to have a potent anti-leukemic effect in non-DS-AMKL and ML-DS, however, consistent drug synergy was only observed in the context of ML-DS. Interestingly, synergistic effects seemed to depend on *JAK* mutational status. The ML-DS patient samples used here harbor different *JAK* mutations, thereby reflecting the mutational landscape of ML-DS – which is characterized by activating mutations in JAKs or other cytokine receptors (Supplemental Figure S3). As ruxolitinib is a JAK1 and JAK2 inhibitor, it exerted synergistically anti-leukemic effects in the *JAK* wild-type and *JAK1* mutated ML-DS samples, while having only additive to mild synergistic effects in samples, which harbor an activating *JAK3* mutation and therefore likely sustain aberrant JAK-STAT signaling after ruxolitinib treatment. Thus, *JAK* mutational status must be taken into account when pursuing further pre-clinical and clinical testing of this drug combination for ML-DS patients. Patients with *JAK3* mutations may potentially benefit from combining a LSD1 inhibitor with a potent JAK3 inhibitor such as Tofacitinib, which is currently approved for the treatment of rheumatoid arthritis.^47^

Concerning clinical testing, there are a number of ongoing clinical trials using different irreversible LSD1 inhibitors in various tumor entities.^26, 27^ Focusing on AML, initial results have already proved the feasibility of LSD1 inhibition as a monotherapy or in combination with retinoic acid – an approach that achieved reduction of leukemic burden while providing a good risk profile in the difficult to treat subgroup of adult patients with relapsed or refractory AML.^24, 25^ In contrast, other clinical trials testing the irreversible LSD1 inhibitor GSK2879552 in AML and myelodysplastic syndrome were terminated mainly because of a high number of adverse events.^26, 27^ These first clinical results indicate that LSD1 inhibition as monotherapy may be potent for AML treatment, but its combination with another targeted therapy could maximize anti-leukemic effects due to drug synergy – just as we have demonstrated with ruxolitinib – while providing a beneficial toxicity profile as a result of the lower drug doses needed to control leukemic growth.

Our study also provides insight into the molecular mechanisms underlying the synergy between LSD1 inhibition and JAK-STAT signaling blockade. As ML-DS blasts already thrive on aberrant JAK-STAT signaling, we demonstrated that pre-treatment with LSD1 broadly activates cytokine signaling which likely contributes to the myeloid differentiation we observed. This mechanism could also be key for sensitizing ML-DS blasts to ruxolitinib, as these cells become even more dependent on constitutively active JAK-STAT signaling – a stimulus which is then abruptly revoked through ruxolitinib treatment. However, the enhanced cytokine signaling observed after LSD1 inhibition should be viewed not only in the context of differentiation signal cascades, but also as an integral part of immunological processes, since we observed upregulation of other immunological gene signatures upon LSD1 inhibition alone or in combination. In the context of solid tumors, LSD1 inhibition has already been demonstrated to increase anti-tumor T cell immunity by inducing interferon signaling and increasing tumor immunogenicity.^48^

Taken together, we are the first to demonstrate synergistic anti-leukemic effects using the combination of an irreversible LSD1 inhibitor and the JAK1 and JAK2 inhibitor ruxolitinib in ML-DS, with both drugs being specifically selected to target core molecular features of the leukemic transformation from TAM to ML-DS.^8^ As the LSD1 inhibitor T-3775440 already displays potent myeloid differentiation and proliferation arrest as monotherapy, adding ruxolitinib drastically increased apoptosis and yielded significant reduction of leukemic burden *in vivo*. However, JAK mutational status has to be considered when combining both drugs, since synergy was only observed in the absence of an activating *JAK3* mutation.

## Author contributions

JG performed experiments, analyzed and interpreted the data and wrote the manuscript. RB performed experiments, analyzed and interpreted the data and revised the manuscript. LG performed experiments. DH and JHK designed the study, analyzed and interpreted the data, wrote the manuscript and academically drove the project.

## Acknowledgements

The authors thank Michelle Ng for assisting in the manuscript preparation. We thank Lonneke Verboon, Hasan Issa, and Oriol Alejo-Valle for their technical support. This work was supported by funding to JHK from the German Federal Ministry of Education and Research (BMBF; MyPred 01GM1911A), the “Hilfe für krebskranke Kinder Frankfurt e.V.”, and the European Research Council (ERC) under the European Union’s Horizon 2020 research and innovation programme (grant agreement #714226). JHK. is a recipient of the St. Baldrick’s Robert J. Arceci Innovation Award.

