## Supplemental Material for "Combining LSD1 and JAK-STAT inhibition targets Down syndrome-associated myeloid leukemia at its core"

Grimm et al.

### Supplemental Tables

**Supplemental Table 1.** Clinical characteristics of patient samples used in this study.

| | gender | age at diagnosis (months) | WBC ($10^9$/L) | hemoglobin (g/dl) | bone marrow blasts (%) | CNS involvement | allogeneic stem cell transplantation | molecular genetics | cytogenetics | response | relapse |
|---|---|---|---|---|---|---|---|---|---|---|---|
| ML-DS #1 | female | 13 | 168.0 | 7.1 | 16.0 | no | no | $NRAS^{mut}$ | NA | CCR | no |
| ML-DS #2 | male | 16 | 4.8 | 12.0 | 10.5 | no | no | $GATA1^{mut}$ | NA | CCR | no |
| ML-DS #3 | male | 26 | 32.5 | 7.9 | 70.5 | no | yes | $GATA1^{mut}$ | 47,XY,t(3;13)(q?26;q?13~14)del(13)(q?14q22),+21c[cp14]/47,sl,del?(15)(q?)[cp2]/46,XY[1] | NR, CCR after allogeneic SCT | no |
| non-DS-AMKL #1 | male | 12 | 40.0 | 9.9 | 64.0 | no | no | $KMT2A^{mut}$ | 46,XY[15].nuc ish 3q26(EVI1x2)[100/100],8q22(RUNX1T1x2),21q22(RUNX1x2)[98/100],11q23(MLLx2)[99/100], 16q22(CBFBx2)[100/100] 17q21.1(RARAx2)[100/100] | CCR | no |
| non-DS-AMKL #2 | male | 2 | 33.8 | 9.5 | 34.0 | yes | yes | NA | 46,XY,t(1;8;22)(p13;q22;q13)[14]/46,XY[1] | NR | yes |

### Supplemental Figures

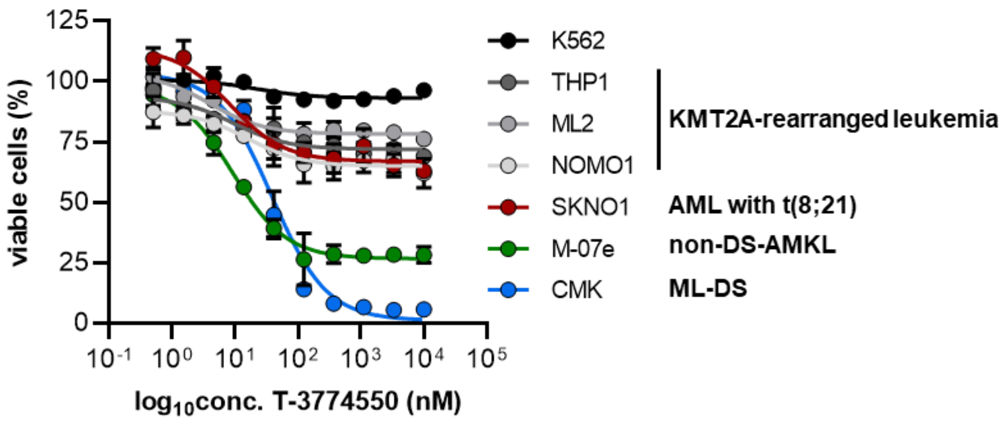


**Supplemental Figure S1. ML-DS and non-DS-AMKL cell lines are more responsive to the LSD1 inhibitor T-3775440 than other pediatric AML subtypes.** Dose-response curves depicting cell viability of different pediatric AML cell lines after treatment with serial dilutions of the LSD1 inhibitor T-3775440 for six days. All cell viability values were normalized to the corresponding DMSO control.

ML-DS, myeloid leukemia associated with Down syndrome; non-DS-AMKL, acute megakaryoblastic leukemia not associated with Down syndrome; AML, acute myeloid leukemia; DMSO, dimethyl sulfoxide.

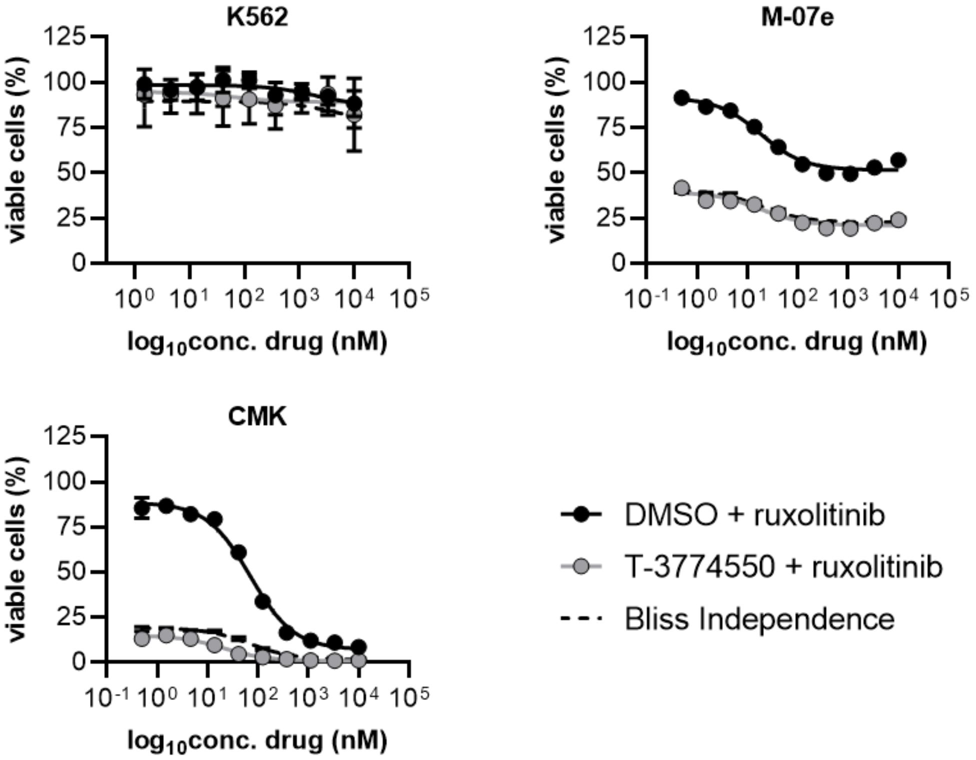


**Supplemental Figure S2. The combination of the LSD1 inhibitor T-3775440 and ruxolitinib exerts synergistic cytotoxic effects in the non-DS-AMKL cell line M-07e and the ML-DS cell line CMK.** Dose-response curves depicting cell viability of different pediatric AML cell lines after treatment with DMSO or 350nM of T-3775440 for six days and the addition of serial dilutions of ruxolitinib from day 3 to day 6. All cell viability values were normalized to the corresponding all DMSO control.

Non-DS-AMKL, acute megakaryoblastic leukemia not associated with Down syndrome; ML-DS, myeloid leukemia associated with Down syndrome; AML, acute myeloid leukemia; DMSO, dimethyl sulfoxide.

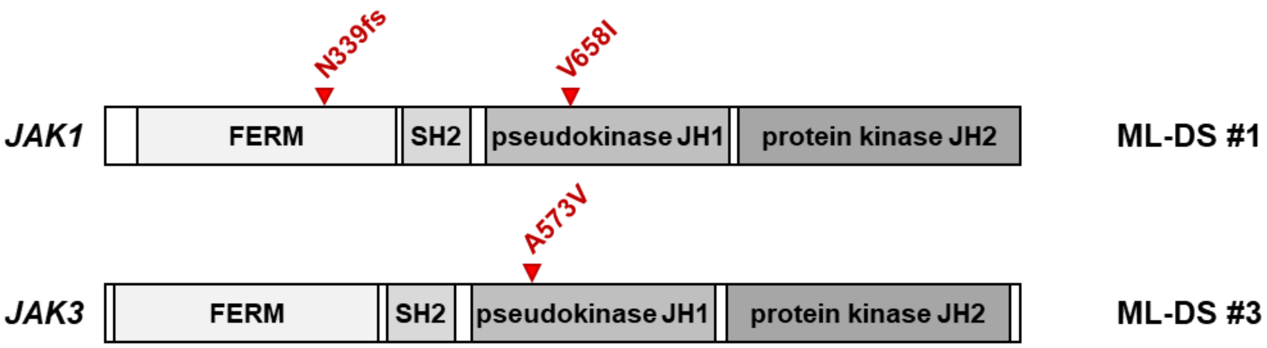


**Supplemental Figure S3. Schematic of the Janus kinases 1 and 3.** Depiction of the different Janus kinase domains and the locations of mutations identified in the ML-DS #1 and #3 patient samples. Adapted from Labuhn et al.

ML-DS, myeloid leukemia associated with Down syndrome.

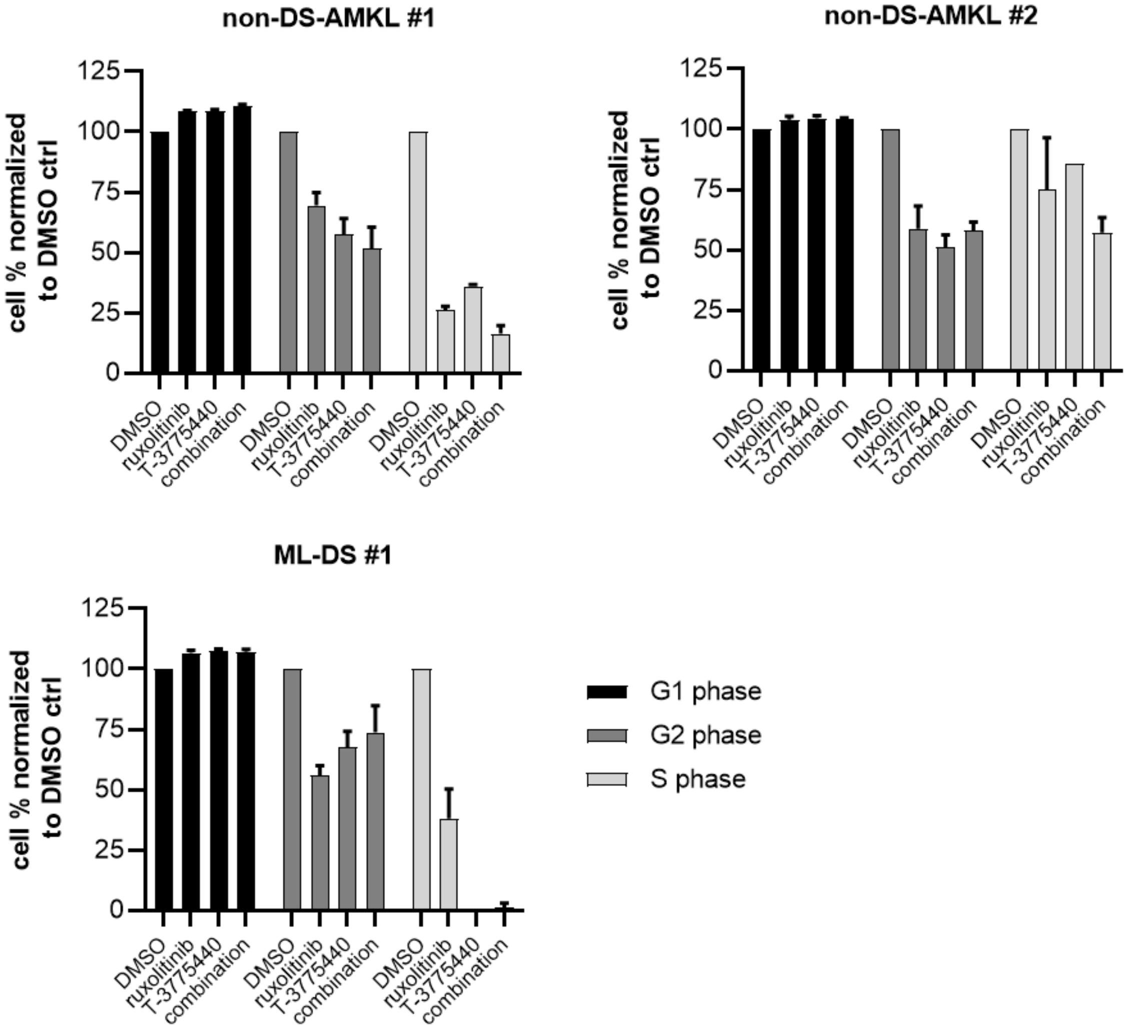


**Supplemental Figure S4. LSD1 inhibitor T-3775440 blocks progression from G1 to S phase in ML-DS and non-DS-AMKL patient samples.** Bar plots displaying the percentage of cells in each phase of cell cycle, as quantified via BrdU assay. All values are normalized to the corresponding DMSO control.

ML-DS, myeloid leukemia associated with Down syndrome; non-DS-AMKL, acute megakaryoblastic leukemia not associated with Down syndrome; BrdU, Bromodeoxyuridine; DMSO, dimethyl sulfoxide.

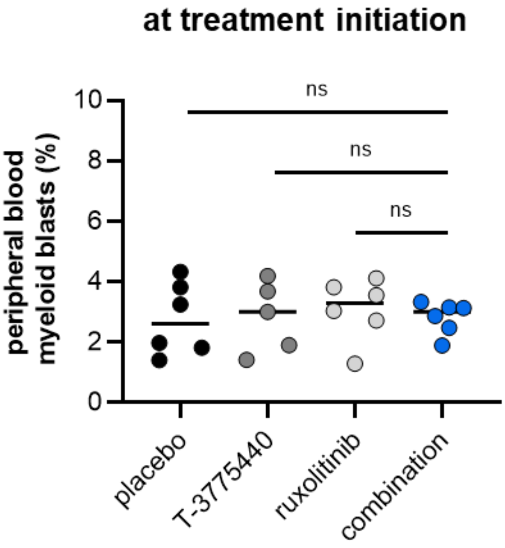


**Supplemental Figure S5. Stable peripheral blood engraftment *in vivo* at the point of treatment initiation.** Percentage of human myeloid blasts in the peripheral blood of humanized immunodeficient mice transplanted with the ML-DS #1 patient sample at the starting point of the treatment period. Groups treated with DMSO, T-3775440, ruxolitinib, or the combination of both drugs are shown separately. ns $P>0.05$; $P$ values are derived from two-tailed Student’s t-tests comparing two groups.

ML-DS, myeloid leukemia associated with Down syndrome.

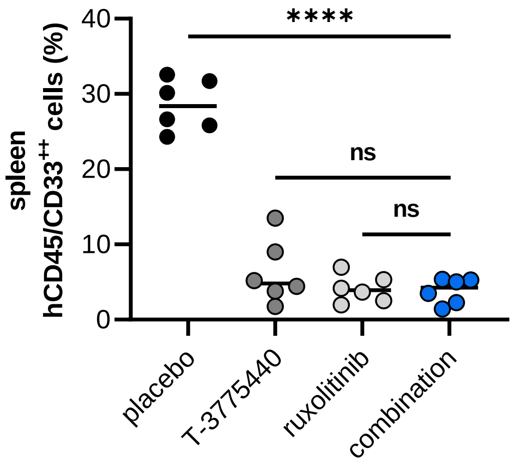


**Supplemental Figure S6. LSD1 inhibitor T-3775440 or ruxolitinib monotherapy, or the combination of both drugs, all significantly reduce leukemic infiltration in the spleens of transplanted mice.** Percentage of human myeloid blasts in the spleens of humanized immunodeficient mice transplanted with the ML-DS #1 patient sample, after seven days of treatment with placebo, T-3775440, ruxolitinib, or a combination of both drugs. ns P>0.05, ****$P$<0.0001; $P$ values are derived from two-tailed Student's t-tests comparing two groups.

ML-DS, myeloid leukemia associated with Down syndrome.

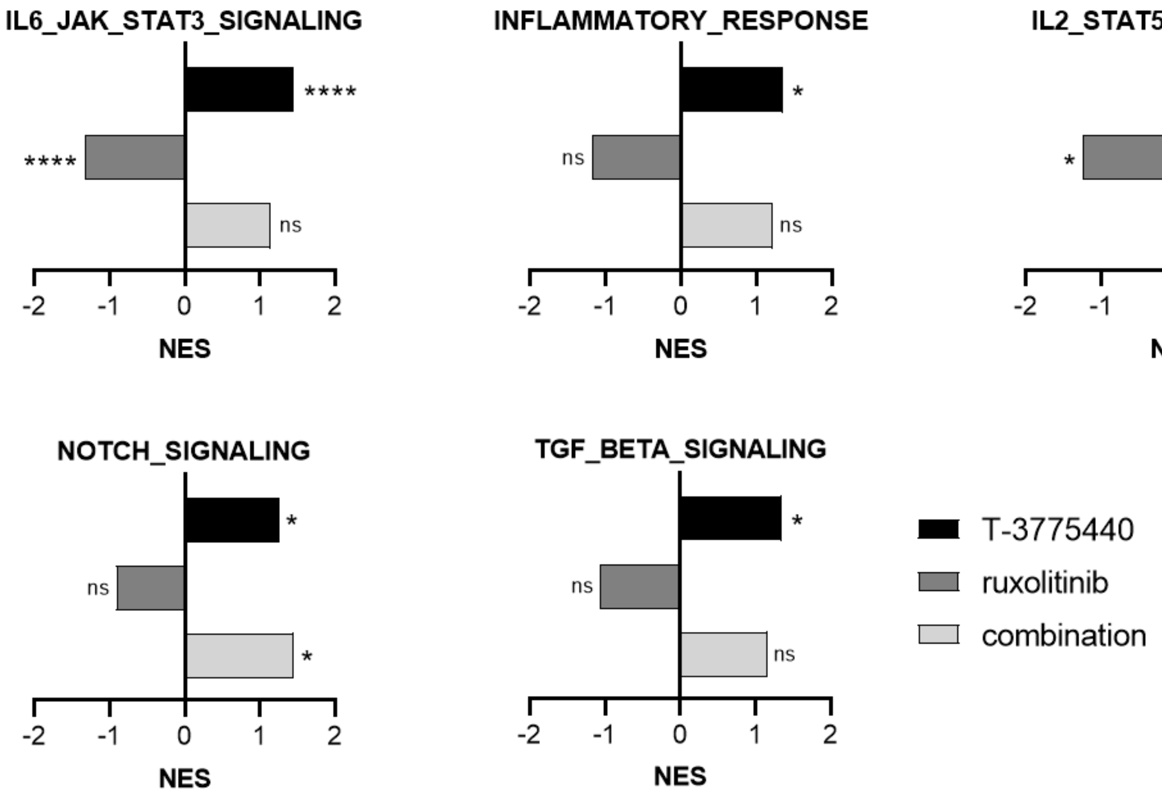


**Supplemental Figure S7. GSEA reveals upregulation of cytokine signaling and pathways involved in differentiation after treatment of ML-DS patient samples with the LSD1 inhibitor T-3775440.** Bar plots depicting normalized enrichment scores (NES) for the respective pathways, as determined using GSEA. This analysis was performed using RNA sequencing data from the patient samples ML-DS #1 and #2 after two days of treatment with DMSO, T-3775440, ruxolitinib, or a combination of both drugs. The DMSO treated cells were used as controls for GSEA. ns *P*>0.05, * *P*<0.05, **** *P*<0.0001; *P* values are calculated by the GSEA software.

GSEA, gene set enrichment analysis; ML-DS, myeloid leukemia associated with Down syndrome; NES, normalized enrichment score; DMSO, dimethyl sulfoxide.
